# Reactivation of TAp73 tumor suppressor by protoporphyrin IX, a metabolite of aminolevulinic acid, induces apoptosis in *TP*53-deficient cancer cells

**DOI:** 10.1101/357293

**Authors:** Alicja Sznarkowska, Anna Kostecka, Anna Kawiak, Pilar Acedo, Mattia Lion, Alberto Inga, Joanna Zawacka-Pankau

## Abstract

**Background:** The p73 protein is a tumor suppressor that shares structural and functional similarity with p53. p73 is expressed in two major isoforms; the TA isoform that interacts with p53 pathway, thus acting as tumor suppressor and the N-terminal truncated ΔN isoform that inhibits TAp73 and p53 and thus, acts as an oncogene.

**Results:** By employing a drug repurposing approach, we found that protoporphyrin IX (PpIX), a metabolite of aminolevulinic acid (ALA) applied in photodynamic therapy of cancer, stabilizes TAp73 and activates TAp73-dependent apoptosis in cancer cells lacking p53. The mechanism of TAp73 activation is *via* disruption of TAp73/MDM2 and TAp73/MDMX interactions and inhibition of TAp73 degradation by ubiquitin ligase Itch.

**Conclusion:** Our findings may in future contribute to the successful repurposing of PpIX into clinical practice.

## Background

Drug repurposing brings hope for the improved treatment of cancer patients due to the financial toxicity of current cancer care that impacts both, patient care and healthcare system worldwide[1].

p73 and p63 are structural and functional homologs of the p53 tumor suppressor protein. All p53 family proteins share the ability to activate p53 target genes involved in apoptosis and cell cycle control under stress conditions.

The *TP*73 gene structure is complex due to the existence of two alternative promoters (P1 and P2). The P1 promoter drives the expression of transcriptionally active TAp73 isoforms, acting as tumor suppressors, while expression from P2 promoter generates N-terminal truncated ΔNp73 isoforms, acting as dominant negative towards TA isoforms and p53 proteins, and thus known as oncogenes[2]. It has been shown that TA and ΔN isoforms possess distinct, often opposing functions in healthy and cancerous tissues[3]. Next, it has been demonstrated that the outcome of chemotherapy depends on the TA and ΔN isoforms ratio[4].

Unlike *TP*53, the gene encoding *TP*73 is rarely mutated in cancers and the functional isoforms are expressed in the majority of human tumors. TAp73 is found inactivated in cancers by binding to oncogenic ΔNp73, MDM2 and MDMX proteins or by degradation by the ubiquitin ligase Itch[5] [6]. Previous studies point at the possibility of exploiting TAp73 as a tumor suppressor for improved cancer therapy. For example it has been demonstrated that p73 *via* activation of c-Jun N-terminal Kinase (JNK) drives the sensitivity to cisplatin in ovarian cancer cells independent on the p53 status[7]. Next, Nutlin3, a selective inhibitor of MDM2, stabilizes TAp73 and induces TAp73-mediated apoptosis[8]. It has also been demonstrated that simultaneous induction of proteotoxic and oxidative stress leads to JNK-induced phosphorylation of TAp73 and TAp73-mediated cell death in p53-null tumors[9].

Small molecule protoporphyrin IX (PpIX), is a natural metabolite of δ - aminolevulinic acid, a pro-drug applied in clinics in photodynamic therapy of cancer (PDT)[10]. PpIX induces HeLa cells’ apoptosis *per se*, without light excitation[11], stabilizes and activates wild-type p53 in human colon carcinoma cells[12] and binds to p73[13].

Here, we have found that PpIX activates TAp73 in cancer cells lacking *TP*53. We demonstrated that PpIX-activated TAp73 compensates for p53 loss in cancer cells and induces apoptosis. The mechanism of transcriptional activation of TAp73 by PpIX is *via* inhibition of TAp73/MDM2 and TAp73/MDMX interactions. TAp73 protein stabilization is achieved by disrupting TAp73/Itch complex by PpIX.

## Results

### Protoporphyrin IX inhibits proliferation and induces apoptosis in *TP*53-null cancer cells

It has been demonstrated that PpIX *per se*, activates wild-type p53 by disrupting p53/MDM2 complex and induces p53-dependent and independent apoptosis in human colon cancer cells[12]. Since PpIX binds to p73[13] we reasoned that p73 might play role in mediating cell death in p53-null cancer cells upon PpIX treatment. Here, in long-term proliferation assay, we showed that PpIX inhibits growth of several *TP*53-null cancer cells in a dose-dependent way (Figure 1a). Interestingly, overexpression of TAp73α sensitized cancer cells to PpIX as shown in short- and long-term proliferation assays (Figure 1b,c). Thus accumulated TAp73 contributed to cancer cells’ susceptibility to PpIX.

**Figure 1.**
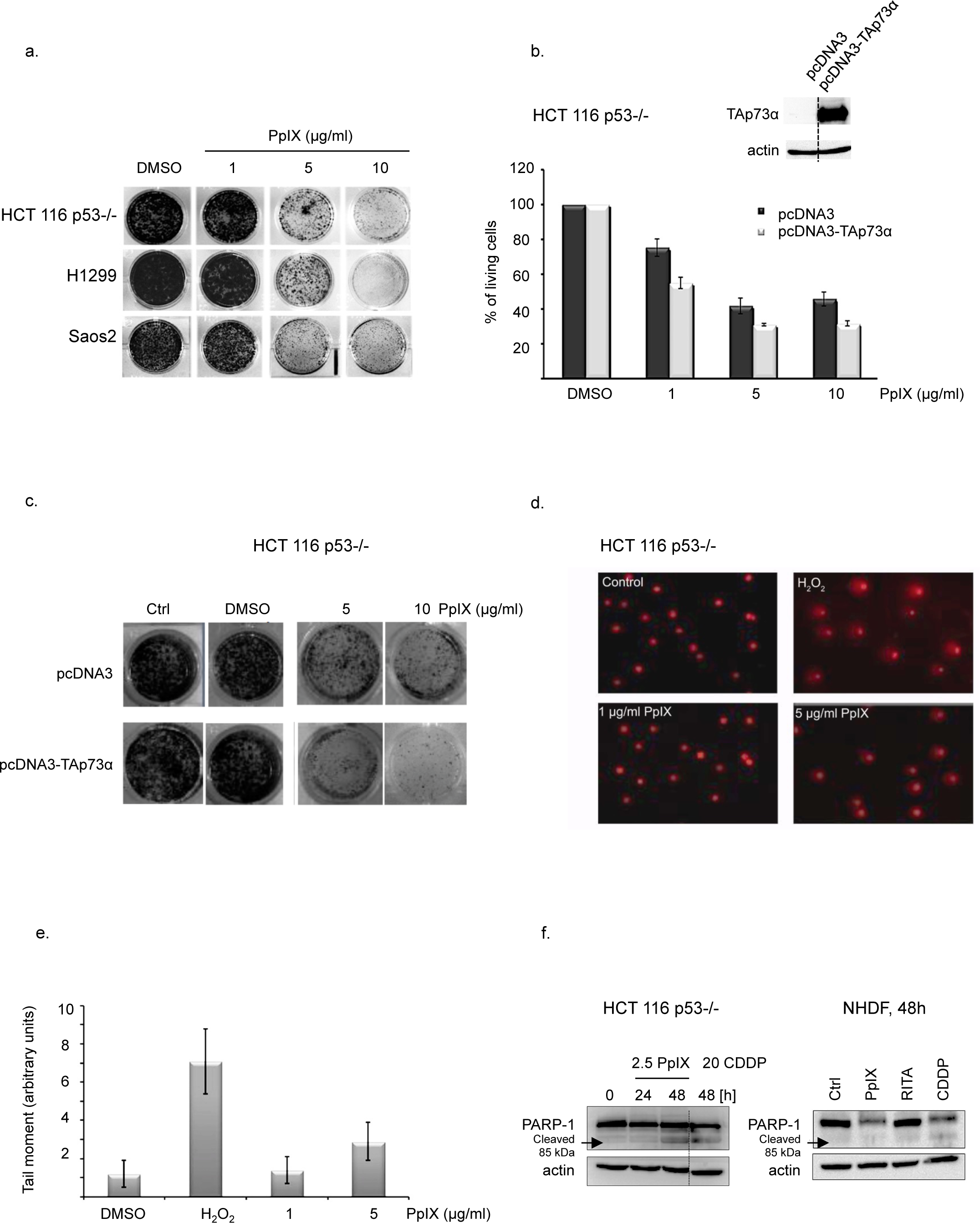
PpIX inhibits proliferation of cancer cells lacking p53. (a) PpIX induces dose-dependent growth inhibition in a long-term proliferation assay. (b) Ectopic expression of TAp73α sensitizes cells to PpIX after 24 h as demonstrated by WST-1 proliferation assay. Inserted blot represents the level of expression of TAp73α. Please note that the blot has been cropped. Dotted line represents where the blot has been cut. The uncropped full length version is pesented in Suppl. Fig. 3a. (c) TAp73α overexpression sensitizes H1299 to PpIX-induced inhibition of proliferation. (d, e) PpIX does not induce DNA damage in cancer cells at effective therapeutic concentrations. (f) PpIX induces PARP-1 cleavage in HCT 116 p53-/- but not in non-transformed human diploid fibroblasts. Dotted line represents where the blot has been cut. The uncropped blot is presented in Suppl. Fig. 3b.

Next, we assessed the genotoxicity of PpIX. The comet assay showed that PpIX did not induce DNA damage in human colon cancer cells (Figure 1d,e) at effective concentrations. Importantly, western blotting demonstrated PARP-1 cleavage, indicating active apoptosis, only in cancer cells but not in normal human diploid fibroblasts (NHDF) after protoporphyrin IX treatment (Figure 1f). A slight reduction in total PARP-1 was detected in NHDF after PpIX and cisplatin (CDDP) but only CDDP induced PARP-1 cleavage in these cells. This implies that PpIX is non-genotoxic and does not affect normal cells at concentrations tested in comparison to cisplatin.

### Protoporphyrin IX induces TAp73 and its apoptotic target genes in *TP*53-null cancer cells

To assess the potential of PpIX against other *TP*53-null human cancer cells than HCT 116 p53-/-, we employed human lung adenocarcinoma cells (H1299) and human osteosarcoma cells (Saos2). The caspase assay showed potent induction of apoptosis by PpIX after 6h in all three cell lines tested (Figure 2a and Supplementary Figure 1a). Induction of active caspases by PpIX correlated with the accumulation of cleaved PARP-1 in H1299 (Figure 2b).

**Figure 2.**
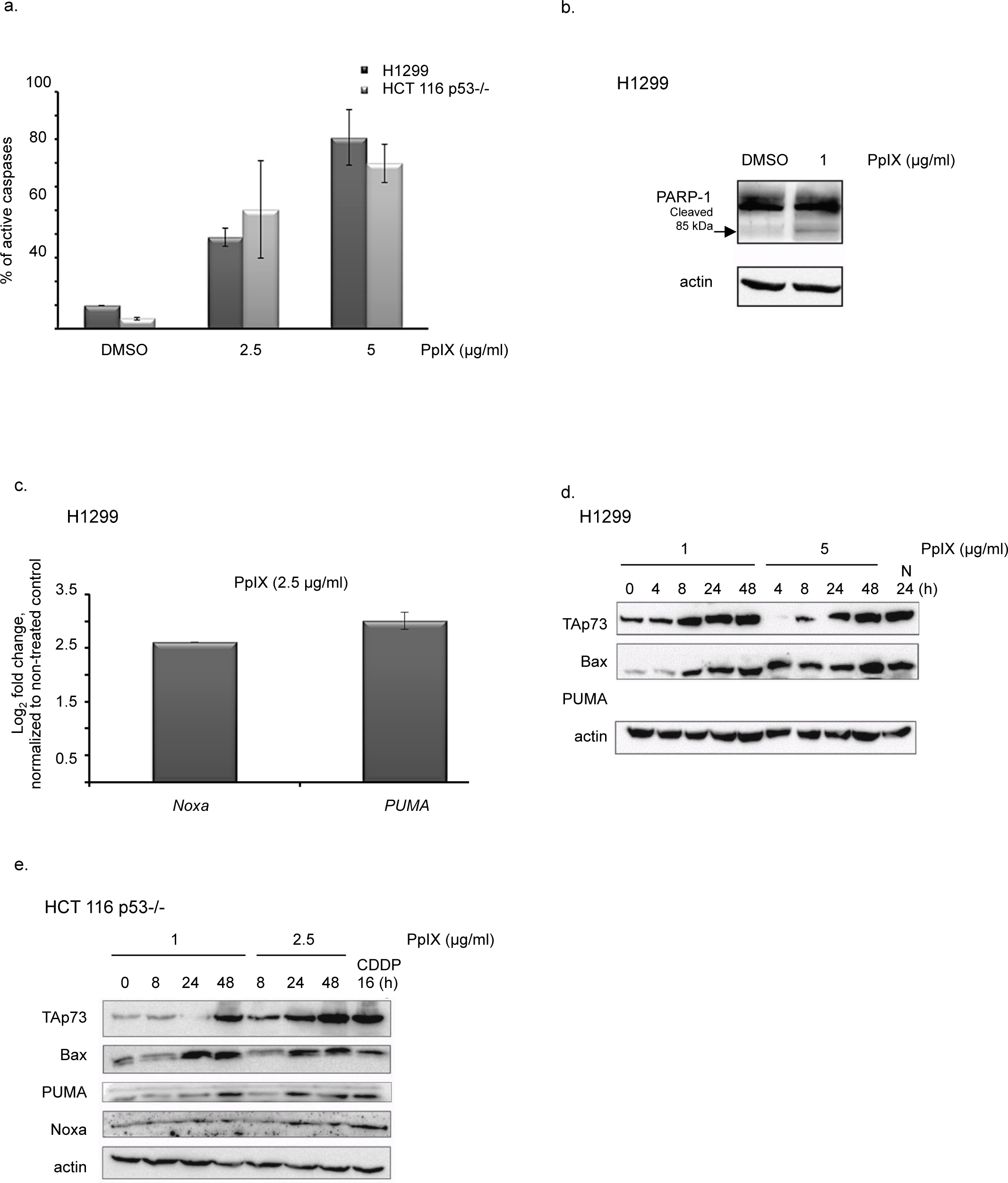
PpIX induces apoptosis and activates TAp73 in p53-null cancer cells. (a) PpIX induces caspases as shown by the increase in the fluorescent signal in p53-null HCT 116 p53-/- and H1299 cancer cells. (b) PARP-1 cleavage in H1299 cells is induced by PpIX after 48 h treatment. (c) mRNA levels of *NOXA* and *PUMA* in H1299 cells treated are elevated with 1 μg/ml PpIX after 12h. (d, e) PpIX induces TAp73 and its proapoptotic targets in H1299 (d) and HCT 116 *TP*53 -/- (e) cancer cells. 20 µM CDDP and 10 µM Nutlin were used as positive controls for induction of p73. The uncropped blots for figure (d) are presented in Suppl. Fig. 4a.

TAp73 and p53 recognize the same target genes involved in the apoptotic response. We found that PpIX at the concentrations from 1 to 5 μg/ml induces accumulation of TAp73 with the concomitant upregulation of its apoptotic targets *Noxa* and *PUMA* on both mRNA and protein levels (Figure 2c,d,e). At the same time we observed the downregulation of the oncogenic ΔN isoform of p73 (data not shown). The induction of TAp73 and its pro-apoptotic targets was also detected for Nutlin3 and cisplatin (Figure 2d and 2e, respectively). Taken together, PpIX stabilizes TAp73 and induces apoptosis in cancer cells lacking *TP*53 in dose- and time-dependent manner.

### PpIX-activated TAp73 compensates for p53 loss in inducing apoptosis in *TP*53-null cancer cells

To determine if TAp73 is critical for PpIX-induced cancer cells’ death, we depleted TAp73 using isoform specific siRNA. PpIX had no effect on the mRNA levels of TAp73 upon the knockdown as demonstrated by real-time PCR (Figure 3a).

**Figure 3.**
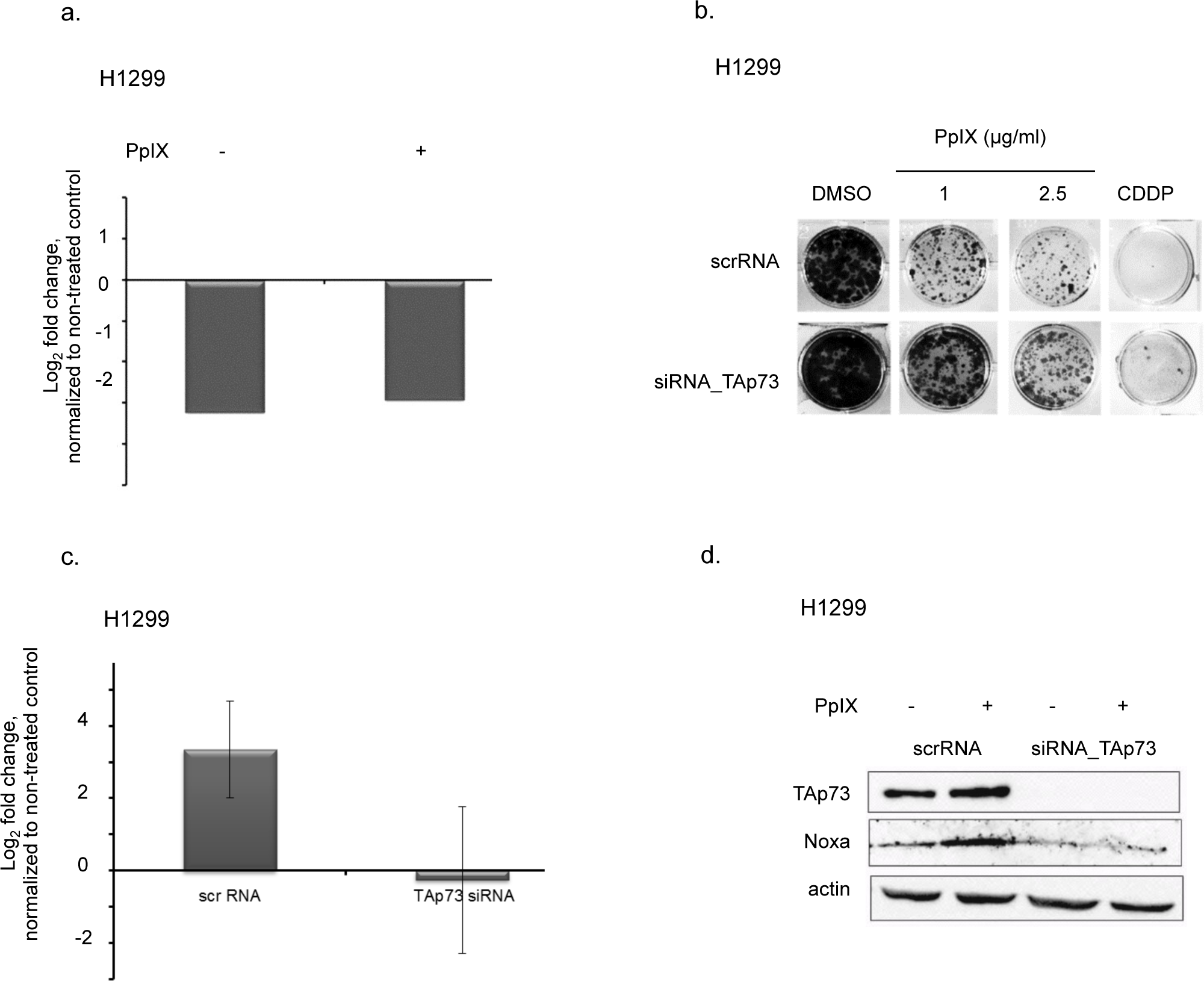
TAp73 knockdown protects cancer cells from PpIX-induced cell death. (a) TAp73 knockdown with siRNA in H1299 cells as confirmed by qPCR. mRNA levels of TAp73 are not affected by PpIX. (b) TAp73 knockdown protects from PpIX-induced growth inhibition in H1299 cells. (c, d) Ablation of TAp73 protects from induction of proapoptotic target Noxa by PpIX on mRNA (c) and protein levels (d).

The knockdown of TAp73 led to resistance to PpIX as shown in the long-term proliferation assay (Figure 3b) and caspase assay (Supplementary Figure 2a). Next, we showed that silencing of TAp73 abolished PpIX-mediated induction of Noxa and Puma as well as PARP-1 cleavage (Figure 3c, d and Supplementary Figure 2b). In addition, the knockdown of TAp73 protected from induction of PARP-1 after treatment of H1299 cells with cisplatin (Supplementary Figure 2c). Thus, our data demonstrated that PpIX-activated TAp73 compensates for p53 loss and induces cancer cells’ death in the absence of p53.

### Protoporphyrin IX activates TAp73 through the disruption of TAp73/MDM2 and TAp73/MDMX complex and protein stabilization

Our results showed that PpIX induces TAp73 transcriptional activity. In cancer cells, p73 transcriptional activity is abrogated, among other mechanisms, by binding to MDM2 or MDMX (human MDM4)[14]. We have shown previously that PpIX inhibits p53/MDM2 complex by binding to the N-terminal domain of p53[12]. Since PpIX also binds to p73[13] we addressed the question if PpIX abrogates interactions between TAp73 and MDM2 by using a defined yeast-based reporter system[15]. Briefly, we expressed TAp73 alone or together with MDM2 or MDM4 in a yeast strain harboring a chromosomally integrated luciferase reporter containing the p53 response element derived from the *PUMA* promoter. Our data manifested that PpIX restored the p73-dependent reporter in yeast strains expressing TAp73/MDM2 or p73/MDM4. This indicated that PpIX ablates TAp73/MDM2 and TAp73/MDM4 interactions and promotes TAp73 transactivation function (Figure 4a). To investigate if PpIX can inhibit TAp73/MDM2(X) interactions also in cancer cells, we immunoprecipated TAp73 after treatment with PpIX and blotted the membrane with MDM2 or MDMX antibodies. Western blot showed inhibition of TAp73/MDM2 and p73/MDMX interactions by PpIX in H1299 and HCT 116 TP53-/-cells (Figure 4b,c).

**Figure 4.**
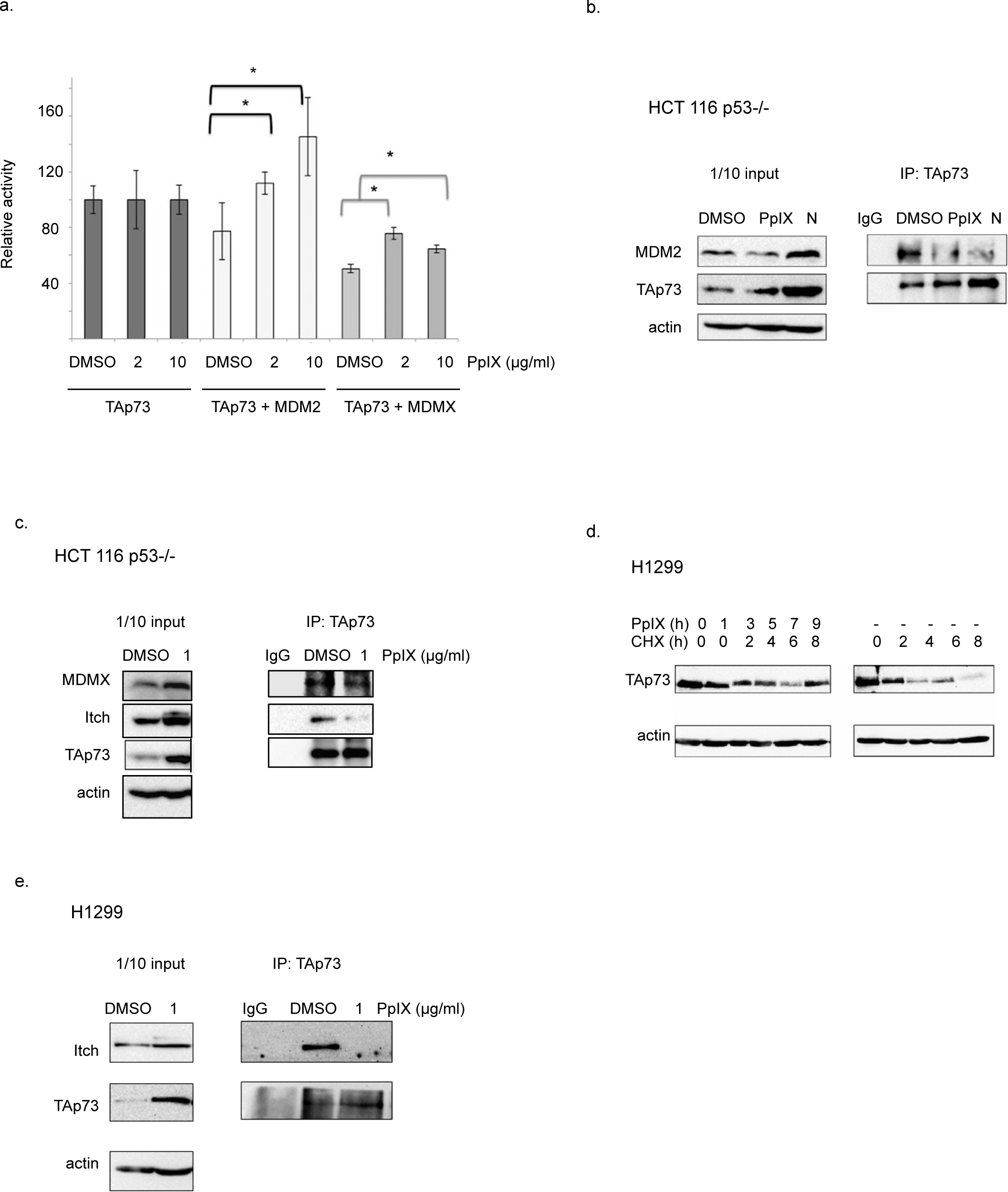
PpIX ablates TAp73/MDM2, TAp73/MDMX and TAp73/Itch complexes. (a) PpIX rescues transcriptional activity of TAp73 thorough ablation of TAp73/MDM2 and TAp73/MDM4 interactions as assessed by yeast-based reporter system. The *t*-student test was performed for statistical analysis with p ≤ 0.05. * samples are considered statistically significant. (b, c) Disruption of TAp73/MDM2 (b) and TAp73/MDMX and TAp73/Itch (c) binding by PpIX is shown in co-immunoprecipitation experiment in HCT 116 p53-/-cells. Uncropped blots are presented in Suppl. Fig. 5a and 5b. N-Nutlin (d) Chase experiment demonstrates stabilization of TAp73 by 1 µg/ml PpIX in H1299. NT-not treated control (e) TAp73/Itch interaction is inhibited by PpIX in H1299 by 1 µg/ml PpIX. The uncropped blots are shown in Suppl. Fig. S6a.

These findings are unexpected, since dual inhibitors of TAp73/MDM2 and TAp73/MDMX interactions have not been described to date.

Since TAp73 protein levels (but not mRNA levels) were upregulated by PpIX in cancer cells (Figure 2d,e), we performed pulse-chase experiments with cycloheximide (CHX) to assess whether PpIX increases the half-life of TAp73 in cancer cells. We observed that PpIX decreased the TAp73 cellular turnover (Figure 4d).

The key E3 ubiquitin ligase responsible for TAp73 degradation is Itch[6]. Small molecules inhibiting Itch activity in cancer cells have recently been considered to have anticancer potential[16]. We reasoned that PpIX might inhibit TAp73/Itch interactions to induce TAp73 stabilization. We then co-immunoprecipitated TAp73 and demonstrated that PpIX inhibits TAp73/Itch in both H1299 and HCT TP53-/-cell lines and the inhibition was more pronounced in H1299 cells (Figure 4 c,e).

Taken together, our data showed that PpIX activates TAp73-dependent apoptosis at several levels, namely, by promoting its transcriptional activity and protecting from proteasomal degradation.

## Discussion

It has been demonstrated that TAp73 KO mice are prone to carcinogen-induced tumorigenesis and that around 35% of mice cohort develop lung adenocarcinomas[17]. Acute genetic ablation of ∆N isoforms triggers rapid regression of thymic lymphomas developed in p53-null mice[18]. Thus, deletion of ∆*Np*63 or ∆*Np*73 can compensate for p53 tumor suppression in thymic lymphomas, and this occurs due to accumulation of TA isoforms of p63 and p73 and induction of apoptosis[19]. Therefore, accumulated TAp73, similarly to p53, can be considered a promising therapeutic target in tumors deficient or mutant for *TP*53 gene.

Cancer-related deaths are a major health problem encountered today[20]. Therefore, there is an urgent need for the development of effective oncology drugs for tailored treatment of patients suffering from the metastatic disease. The success of anti-cancer treatment relies on the strategy that allows inhibition of cancer drivers without the induction of apparent side effects. However, such strategies are still at early stage of development.

The p53 reactivation suppresses established tumors *in-vivo*[21], yet this strategy is not implemented therapeutically. APR-246/MQ, a compound, in Phase II clinical development (clinical trial ID: NCT03268382), that binds to cysteine residues in mutant p53 and reactivates its function[22] brings hope for patients harboring *TP*53 gene mutations.

Given that approximately around 2/3 of all human cancers harbour *TP*53 gene mutations (https://p53.fr/), a development of a new approach to compensate for p53 loss, is of the outmost clinical relevance. Small molecules reactivating TAp73 are of a special interest. Few molecules directly or indirectly activating TAp73 in cancer cells have been described in literature. Small molecule Nutlin3, a potent inhibitor of p53/MDM2 interactions was found to activate p73 and induce p73-mediated apoptosis by disrupting p73/HDM2 association in cancer cells lacking p53[8]. Next, small molecule RETRA was described to specifically suppress mutant p53-bearing tumor cells *in vitro* and in mouse xenografts by disrupting mtp53/p73 complex[23]. The widely-used genotoxic drug cisplatin (CDDP) was also shown to induce TAp73-mediated apoptosis in ovarian cancer cells irrespective of p53 status. Recent study showed that TAp73 sensitizes p53-null colon cancer cells to bortezomib and TAp73 was shown to compensate for p53-loss in induction of apoptosis after drug treatment[24].

Our previous findings showed that protoporphyrin IX binds to p53 and activates p53-dependent and independent cell death in colon cancer cells[12]. Consistently, PpIX was shown to induce cancer cell death in sarcoma cells[25]. In line with our data (Figure 1d), it was demonstrated that PpIX does not induce DNA damage but induces chromatin condensation by triggering the translocation of factors such as AIF from mitochondria to the nucleus, where it binds to the DNA and provokes caspase-independent chromatin changes[25].

TAp73 bears high structural and functional homology with p53. This prompted us to study if PpIX activates TAp73 in tumors lacking functional p53. Here, we showed that PpIX inhibits the growth of cancer cells lacking p53 in a dose-dependent manner. Fluorescent-based caspase assay indicated that the growth suppression resulted from induction of apoptosis. This is in line with previous study in which we observed inhibition of proliferation and the accumulation of HCT TP53-/-cells in the sub-G1 fraction of the cell cycle after PpIX exposure[12].

Recently, detailed analysis of the oligomeric state of TAp73 alpha by Melino and Dötsch labs revealed a high structure homology between p53 and p73 tetramers[26]. This shows that p73 can be pharmacologically restored in a way similar to p53. This is supported by our previous and current findings, which showed that PpIX binds to p73 and induces accumulation of TAp73 and its apoptotic targets in cancer cells in a fashion similar to p53. Using siRNA we determined that TAp73 is crucial for induction of apoptosis in cancer cells treated with PpIX. This supports the notion that pharmacologically restored TAp73 behaves similarly to activated wild-type p53 and that TAp73 can compensate for p53 loss in cancer cells lacking *TP*53 gene.

To gain better insight into the mechanism of activation of TAp73 in cancer cells by PpIX, we used a yeast-based reporter system previously developed to screen for activators of p53[15]. PpIX restored TAp73 reporter in the presence of MDM2 or MDM4, inhibitors of p73 transcriptional activity. Thus, we concluded that PpIX abrogates TAp73/MDM2 and TAp73/MDMX interactions to induce its transcriptional activation. In agreement with the above, PpIX inhibited TAp73/MDM2 and TAp73/MDMX interactions in cancer cells. This finding is novel since small-molecule, dual inhibitors of MDM2 and MDMX have not been described yet. So far only stapled peptides therapeutic ALRN-6924, targeting both MDM2 and MDMX and stabilizing wild-type p53[27] have been developed and is currently in Phase I clinical trial in wild-type p53 AML and MDS (clinical trial ID: NCT02909972). Of note, our findings have important clinical relevance since the ability to disrupt the interactions of TAp73 with both MDM2 and MDMX is a favorable therapeutic approach in tumors with MDMX amplification.

Since mRNA levels of p73 were not elevated by PpIX we reasoned, that the increase on the protein level must be related to the prolonged stability of TAp73 in cancer cells. Chase experiments confirmed that PpIX stabilizes TAp73 on protein level. The stabilization was a consequence of disruption of interactions between TAp73 and E3 ubiquitin ligase, Itch in cancer cells.

## Conclusions

Taken together, our data showed that PpIX, a metabolite of aminolevulinic acid, activates TAp73 by several mechanisms converging on activation of its transcriptional activity and protein stabilization leading to transactivation of pro-apoptotic *PUMA* and *NOXA*. Next, pharmacologically activated TAp73 compensates for p53 loss and induces apoptosis in cancer cells. Our findings, might in future lead to successful repurposing of porphyrins into clinical application to treat tumors with *TP*53 gene mutations.

## Methods

All experiments including protocols were performed in accordance with the guidelines and regulations of Karolinska Institute and the University of Gdansk.

### Cell culture and chemicals

The p53-deficient human colon cancer cells HCT 116 *TP53* -/-[28] (a generous gift from Bert Vogelstein, Johns Hopkins Oncology Center, Baltimore, MD, USA), human non-small cell lung carcinoma cell line H1299 (ATCC) and osteosarcoma Saos2 (ATCC) cells were cultured at 37°C in a humidified incubator with 5% CO_2_. All cell lines were maintained in Iscove’s modified Dulbecco’s medium (Invitrogene, Sweden) supplemented with 1 mM sodium pyruvate and 10% FBS (Gibco, Sweden).

Protoporphyrin IX was purchased from Sigma-Aldrich, Germany and dissolved in 100% DMSO (Sigma-Aldrich, Germany). The final concentration of DMSO in cell medium was 0.5%. Cisplatin (CDDP) (Sigma-Aldrich, Germany) was dissolved in dH_2_O and used at final concentration 20 μM. Nutlin (Calbiochem, Sweden) was dissolved in DMSO and used at final concentration 10 μM.

### DNA constructs and siRNA

For ectopic expression of TAp73alfa, we used pcDNA3.1-TAp73α construct kindly provided by Prof. Matthias Dobbelstein. The construct has been sequenced to confirm the fidelity of the transgene sequence. Transfection was performed with Lipofectamine™ 2000 (Invitrogen, Sweden) for 24 h according to the manufacturer’s protocol.

The following siRNAs were used: TAp73_318 sense: AGGGCAUGACUACAUCUGU; antisense: ACAGAUGUAGUCAUGCCCU and TAp73_223 sense: ACCAGACAGCACCUACUUC; antisense: GAAGUAGGUGCUGUCUGGU[29]. Transfection with siRNA (10 nM) was performed with HiPerfect reagent (Qiagene, Germany) for 24 h (for WB analysis) or 48 h (for colony formation assay or qPCR analysis) following manufacturer’s instruction.

### Growth assay

For short-term proliferation assay, HCT 116 TP53-/- cells were seeded at 5^10^3^/well in 96-well plate and transfected with either the empty vector or pcDNA3 containing TAP73α sequence. After 24 h cells were treated with PpIX and allowed to grow for another 24 h. Cells’ viability was measured by WST-1 proliferation reagent (Roche, Switzerland).

For a long-term colony formation assay, 3000 cells/well were seeded into twelve-well plates and treated with indicated concentrations of PpIX. The medium was changed after 3 h and cells were allowed to grow for 7 days. The colonies were stained with crystal violet reagent according to standard protocol.

### Quantitative PCR

48 h after siRNA transfection cells were harvested, mRNA was isolated and reverse transcribed to cDNA according to the manufacturer’s instructions (5 Prime, Hamburg, Germany; Invitrogen, Sweden). For qPCR, the following concentrations were used: 150 nM primers; 10 ng cDNA; 7.5 μl 2 × master mix (Bio-Rad, Sweden); water to a total of 15 μl; Primers used: *TAP*73 forward: GGGAATAATGAGGTGGTGGG and *TAP*73 reverse: AGATTGAACTGGGCCATGAC, *NOXA* (PMAIP1) forward AAGTGCAAGTAGCTGGAAG, reverse: TGTCTCCAAATCTCCTGAGT, *PUMA* forward: CTCAACGCACAGTACGAG and reverse: GTCCCATGAGATTGTACAG, *GAPDH* forward: TCATTTCCTGGTATGACAACG and reverse: ATGTGGGCCATGAGGT

### Yeast-based reporter system

The TAp73-dependent yeast reporter strain yLFM-PUMA containing the luciferase cDNA cloned at the *ADE2* locus and expressed under the control of PUMA promoter response element was transfected with pTSG-p73, pLS-MDM2 (derived from pRB254, generously provided by Dr. R. Brachmann, Univ. of California, Irvine, CA, USA)[15], or pLS-Ad-MDM4 and selected on double drop-out media for TRP1 and HIS3. Luciferase activity was measured 16 hrs after the shift to galactose-containing media and the addition of PpIX (Sigma-Aldrich, Germany), or DMSO. Presented are average relative light units and the standard errors obtained from three independent experiments each containing four biological repeats. Treatment with PpIX did not affect the transactivation of the reporter by TAp73 alone.

### Co-Immunoprecipitation and Western blotting

For TAp73/Itch and TAp73/MDMX co-IP cells were treated for 24 h with 1 µg/ml PpIX and 30 μM MG-132 was added 3 hours before cell harvest. For p73/MDM2 co-IP cells were treated with PpIX or Nutlin for 24 h (HCT TP53 -/- cells). Cells for both whole cell lysates and immunoprecipitates were solubilized in lysis buffer: 25 mM Tris HCl, pH 8.0, 150 mM NaCl and 1% Nonidet P-40 (0.5% for co-IP). For co-IP, 1 mg protein was immunoprecipitated with 1.5 μg of α-TAp73 rabbit polyclonal antibody (Bethyl Laboratories, TX, USA) or normal mouse IgG (Millipore, MA, USA). Immuno-complexes were absorbed onto 40 μl of Dynabeads® Protein A (Invitrogen, Sweden) for 5 h at 4°C. The immunoprecipitates were washed with 1 mL of lysis buffer. The antibodies used for detection were: anti-p73 monoclonal antibodies[18] (IMG 246, Imgenex, UK), anti-MDM2 (Santa Cruz, Germany), anti-Itch (Calbiochem, Sweden), anti-MDMX (Bethyl Laboratories, TX, USA).

Western Blot was performed according to the standard protocol. 100 μg of total cell lysate was subjected to electrophoresis and the following antibodies were used to detect proteins: anti-TAp73 (Bethyl Laboratories, TX, USA), anti-Bax (Santa Cruz, Germany) anti-PUMA (Cell Signaling), anti-Noxa (Calbiochem, Sweden), anti-PARP1/2 (Santa Cruz, Germany), anti-actin (Sigma-Aldrich, Germany).

### Caspase activation assay

Activation of caspases by PpIX was measured with FAM-FLICA™ Poly Caspase Assay Kit (ImmunoChemistry Technologies, Germany) according to manufacturer’s protocol. Caspase activation was monitored by the means of flow cytometry with BD FacsCalibur after 6 h treatment with PpIX.

### Protein stability assay

H1299 cells were pre-treated with 1 µg/ml PpIX for 1 h and translation was inhibited by adding cycloheximide (Sigma Aldrich, Germany) to final concentration 30 µg/ml. Cells were harvested 2, 4, 6 and 8 h after addition of CHX and subjected to Western blot analysis.

### Comet assay

HCT TP53 -/- cells were seeded into 24-well plates and treated with PpIX (1 μg/ml and 5 μg/ml), H_2_O_2_ (100 μM) was used as a positive control. Following treatment, cells were pelleted by centrifugation at 1500 rpm and suspended in 60 μl of PBS (pH 7.4). 10 μl of cell suspension was mixed with 100 μl of 1% low melting agarose (Prona, Reducta LM, Poland) and 75 μl of this cell–agarose mixture was spread on microscopic slides pre-coated with 1% agarose. A third layer of 0.5% low melting agarose (75 μl) was applied over the layer of agarose with the cell suspension. Slides were incubated for 1 h in a lysis solution (2.5 M NaCl, 100 mM EDTA, 100 mM Tris, 1% Triton X-100, pH 10). The microscopic slides were immersed in an alkaline buffer (300 mM NaOH, 1 mM disodium EDTA) for 30 min after which they were subjected to electrophoresis at 1V/cm for 15 min. Subsequently slides were neutralized with a neutralization buffer (0.4 M Tris, pH 7.5) for 15 min and stained with ethidium bromide (20 μg/ml). Cells were analyzed under a fluorescence microscope (Nikon PCM-2000) using Cometscore® software. Images of 20 cells from three slides were analyzed. Densities were measured for each image in two areas: the whole cellular DNA and the area containing only the comet head. Results are presented as tail moment, which is the percentage of DNA in the comet tail multiplied by the tail length.

### Statistical analysis

Yeast assay was performed in 3 independent experiments with 3 triplicates. P < 0.05 was considered statistically relevant. The statistical significance was assessed by a parametric Student’s t test (for variances Fisher–Snedecor’s test was applied and the normality was estimated with the Shapiro–Wilk’s test).

## Abbreviations

TA: transcriptionally active
PpIX: protoporphyrin IX
CDDP: cisplatin
CXH: cycloheximide

## Data availability

Not applicable.

## Authors’ contribution

A.S., A.K., A.K., P.A., M.L., prepared figures and drafted the manuscript; J.ZP. and A.I., designed the study, wrote and revised the manuscript. All authors read and approved the final version of the manuscript.

## Acknowledgement

Research was funded by the grant from the Polish National Science Center no 6126/B/P01/2010/38 NN405612638, Karolinska Institute and Stockholm Läns Landsting, Åke Wibergs stiftelse, IARC IG grant (#12869), the Strategic Research Programe in Cancer (StratCan) and ETIUDA grant. JZP would like to acknowledge the award for Young Talented Scientist from Polish Ministry of Science and Higher Education. We thank Prof. G. Selivanova, Karolinska Institute, for scientific discussions. Thanks are addressed to Katarzyna Koczergo for a technical support. We also thank all our colleagues for sharing their reagents.

